# Easy Kinetics: a novel enzyme kinetic characterization software

**DOI:** 10.1101/674051

**Authors:** Gabriele Morabito

## Abstract

Here will be presented the software Easy Kinetics, a publicly available graphical interface that allows rapid evaluation of the main kinetics parameters in an enzyme catalyzed reaction. In contrast to other similar commercial software using algorithms based on non-linear regression models to reach these results, Easy Kinetics is based on a completely different original algorithm, requiring in input the spectrophotometric measurements of ΔAbs/min taken twice at only two different substrate concentrations. The results generated show however a significant concordance with those ones obtained with the most common commercial software used for enzyme kinetics characterization, GraphPad Prism 8©, suggesting that Easy Kinetics can be used for routine tests in enzyme kinetics as an alternative valid software.

## Introduction

The continuous and rapid evolution of modern biochemical methods make the study of enzyme’s kinetic very useful both in academic research, to test how interesting polypeptidic chain’s variation impact on enzymes functionality, and in industrial processes, to optimize the production processes of the molecules of interest in enzymatic reactors [2]. The Michaelis-Mentem reaction mechanism was proposed almost a century ago to describe how the reaction speed of enzymes is affected by the substrate’s concentration [3], and it’s still the core reference model to describe enzymes kinetics. This model however requires a few parameters to fit the raw data: *nH*, K_m_and V_max_. Several methods were developed by biochemists during years to evaluate these parameters from the raw data, the most used of which allow software like GraphPad Prism 8© [1] to apply linear or non-linear regression model [4]. Original alternative methods for K_m_ and V_max_ determination were proposed, which graphically determine these values [5], but like the previous ones they require multiple spectrophotometric measurements of ΔAbs/min (at least 6 conducted in duplicate) at different substrate concentrations to precisely determine the main kinetic parameters. In this paper will be presented an alternative method implemented in the software Easy Kinetics, which allows determination of the main kinetics parameters of an enzyme catalyzed reaction and the corresponding kinetics graphs, by the spectrophotometric measurements of ΔAbs/min taken twice at only two different substrate concentrations.

## Materials and methods

### Algorithm used in evaluation of K_m_ and V_max_

The evaluation of K_m_ and V_max_ by the spectrophotometric measurements of ΔAbs/min taken twice at only two different substrate concentrations, is based on a trigonometric demonstration (**Fig.1**). Briefly the algorithm transforms the mean of the duplicates at the two measurements in their reciprocal values, considering the Lineweaver-Burk reciprocal plot. Known two points of this graph, it’s universally accepted that they can be joined by one and only one straight line. This line will have an unknown inclination “a” and will intersect the Cartesian axes in points -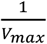 and 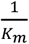, also unknown. However by tracing the projections of the two known points (x_1_,y_1_) and (x_2_,y_2_) on the Cartesian plane, it is evident that the parallel lines y = y_2_ and y = 0 intersect the studied straight line. By the Alternate Interior Angles theorem [6], if two parallel lines are cut by a transversal one, then the pairs of alternate interior angles are congruent: so, by **Fig.1**, “a” = “a_1_”. Considering instead the lines y = y_2_and y = y_1_, which are also parallel and intersected by the studied straight line, for the same theorem discussed before, their internal alternate angles are congruent: so, by **Fig.1**, “a_1_” = “a_2_”. This implies that:

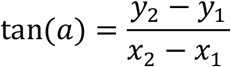

**Fig 1.**
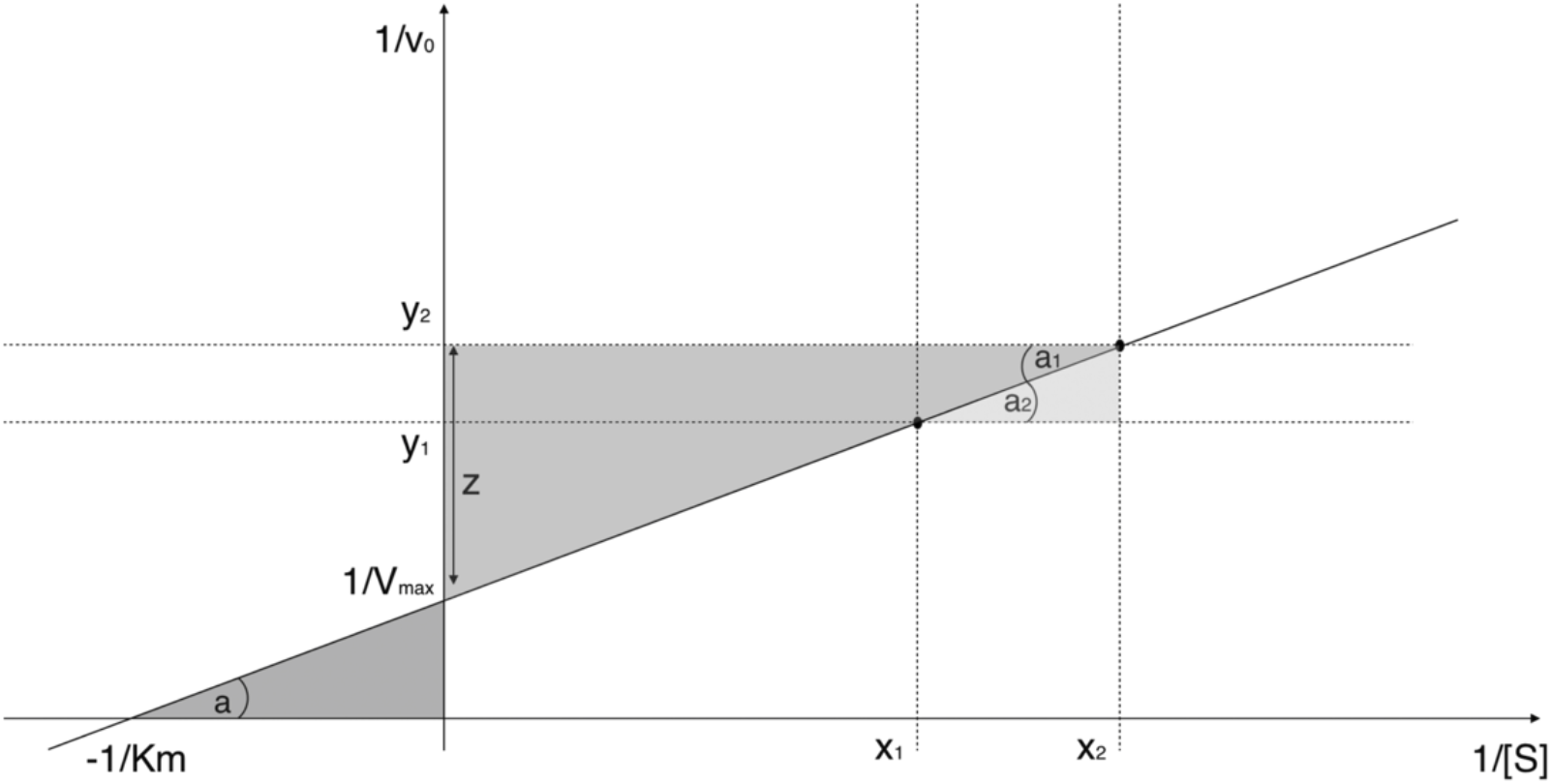
Graphical trigonometric demonstration of the K_m_ and V_max_ evaluation based on Lineweaver-Burk reciprocal plot.

But also 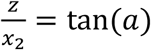, with 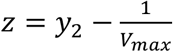, so:

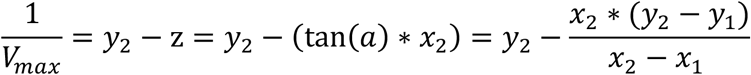

Once calculated 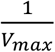, the value of 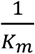 can be determined as follow:

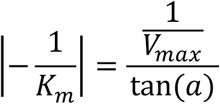

Inverting the two previous values, *K*_*m*_ and *V*_*max*_ will be finally found and from there by subsequent biochemical relations the other kinetic parameters will be deduced (*Supplementary material*) [7-10].

Since Easy Kinetics receives in input only two spectrophotometric measurements, despite being performed twice, if one of these measures is anomalous, it won’t be corrected by other measurements as occurs with regression models; so the software may fall in error. Thus to minimize experimental bias the algorithm is implemented to consider the ΔAbs/min at one substrate concentration only if both the duplicates fall within the range of their mean ± 10% of their average, otherwise the software suggest to repeat these measurements for the substrate concentration considered.

### Software implementation and distribution

Easy Kinetics was developed in C# language with a GPL-3.0 license, both for the versatility of C# and the design object-oriented, for Windows 10 environment (with October update installed), because of the diffusion of Windows 10 and the consequent ease of software distribution. The software installation package can be downloaded freely as windows application on Microsoft Store at the URL:https://www.microsoft.com/it-it/p/easy-kinetics/9nx1f4q5fpg5?activetab=pivot:overviewtab) or alternatively on the repository GitHub (DOI: 10.5281/zenodo.3242785) at the URL:https://github.com/ekin96/EasyKinetics. Easy Kinetics allows the user to operate in 5 different environments several kinetics analyses: “Simple Enzyme Kinetics”, “Inhibition Kinetics”, “Enzymatic Units Assay”, “Calculation of ΔAbs/min”, “Bradford Assay”. Furthermore Easy Kinetics was optimized to self-detect possible substrate-inhibition kinetics.

### Statistical analysis

All statistical analyses were conducted in the software R (v 3.6.1) [11], using both Easy Kinetics and GraphPad Prism 8 [1] for the kinetic analyses. Detailed statistics for all the experiments can be found in the figure legends and/or in the manuscript together with the *n* and definitions of center and dispersion. In all figures, *n* represents the number of different substrates that were used. Before using the Pearson’s or Spearman’s correlation test and using C.V. or Q.C.V. as error value, normality of the variables was checked using the Shapiro-Wilk normality test. Statistical significance was defined for p < 0.05 in all comparisons and calculated as described in the manuscript and/or figure legends.

## Results

### Software interface

Easy kinetics provides to users 5 different user-friendly working environment, “Enzymatic Units Assay” and “Bradford Assay” can both be used alone or included inside “Simple Enzyme Kinetics”, which represent the main environment of the software (**Fig.3**). Both the “Simple Enzyme Kinetics” and “Inhibition Kinetics” environment allow users to generate basal kinetic graphs based on the kinetic parameters evaluated.

**Fig. 2.**
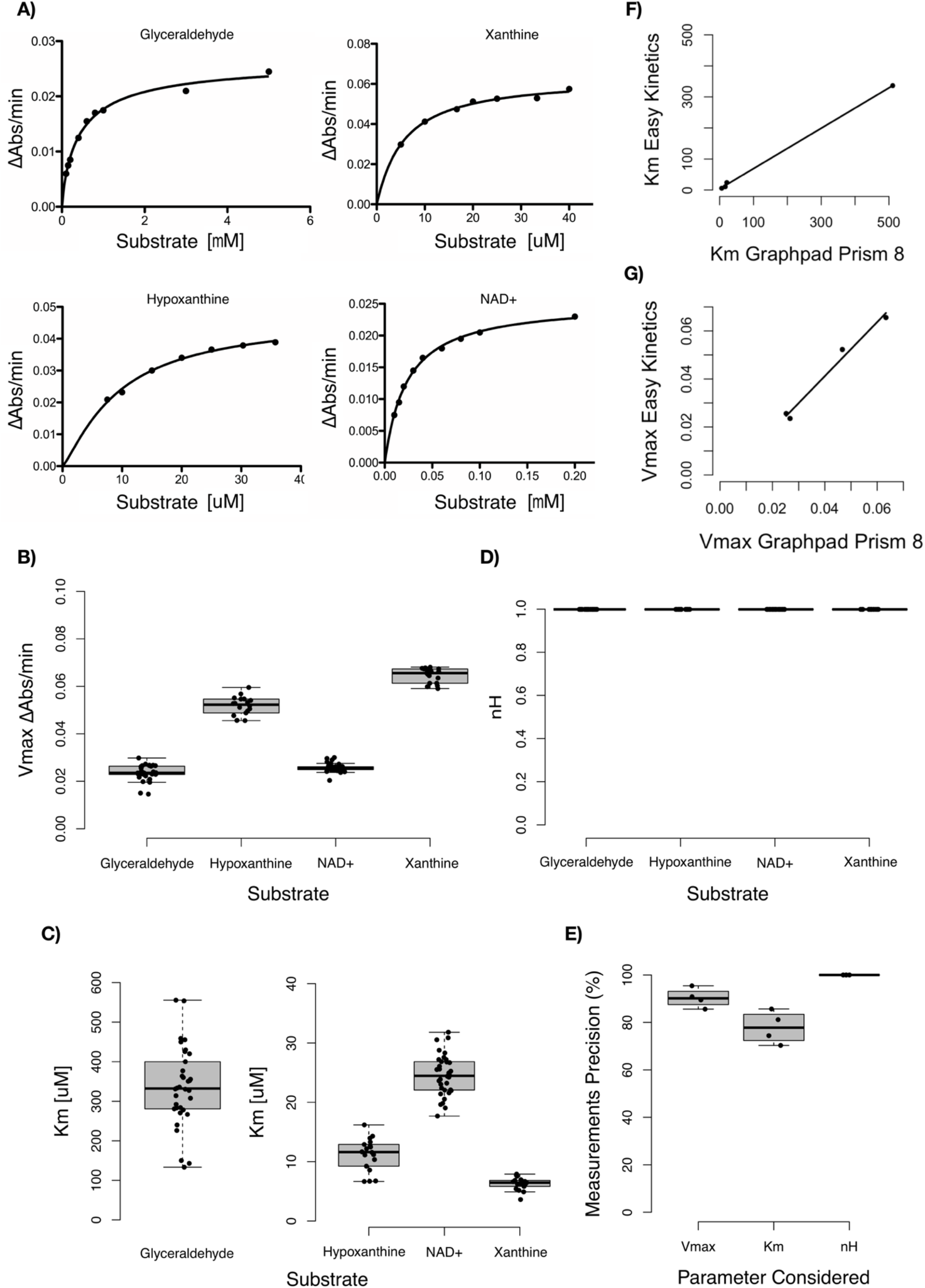
Model’s precision analysis: **A)** Kinetic curves of the two enzymes tested for different limiting substrates, respectively Xanthine or Hypoxanthine for bovine Xanthine oxidase and NAD+ or Glyceraldehyde for bovine Aldehyde dehydrogenase, all the curves were generated using GraphPad Prism 8. **B-D)** Boxplots representing the distributions of Vmax, Km and nH values generated with Easy Kinetics using every two points non repeated permutation of the raw data experimentally obtained. Glyceraldehyde values show a normal distribution for Km (W = 0.96956, p-value = 0.4494) and nH, while a non normal distribution for Vmax (W = 0.89738, p-value = 0.003915). Hypoxanthine values show a normal distribution for Vmax (W = 0.96286, p-value = 0.6576), Km (W = 0.94657, p-value = 0.3738) and nH. NAD+ values show a normal distribution for Km (W = 0.98863, p-value = 0.9728) and nH, while a non normal distribution for Vmax (W = 0.90018, p-value = 0.004639). Xanthine values show a normal distribution for Km (W = 0.95116, p-value = 0.4751) and nH, while a non normal distribution for Vmax (W = 0.88084, p-value = 0.03281). Values outliers in position: 1,7,8,33,38,40,42,51,78,87,105 of the two point non repeated permutations dataset were eliminated for differing too much in Vmax and/or Km estimation from the central value of the distribution. **E)** Boxplots representing the precision of Vmax, Km and nH distribution central value for all the enzymes and limiting substrate tested (n=4), since Km and nH show a normal distribution of values for all the tested substrates, the precision of the estimation was calculated as (1-C.V.)*100, while for Vmax showing non normal distribution for 3 of the tested substrates, the precision was calculated as (1-Q.C.V.)*100. C.V. refers to the coefficient of variation while Q.C.V. to the quartile based coefficient of variation. **F-G)** scatterplots showing graphically the linear correlation between the Vmax and Km generated both by Easy Kinetics (n=4) and GraphPad prism 8 (n=4). The tested substrates show a normal distributions of medians values for Vmax generated using Easy Kinetics (W = 0.87392, p-value = 0.3133) and of values generated using GraphPad Prism8 (W = 0.88607, p-value = 0.3652), allowing use of Pearson’s linear correlation test, while a non normal distribution of mean values for Km generated using Easy Kinetics (W = 0.67175, p-value = 0.005173) and of values generated using GraphPad Prism8 (W = 0.65386, p-value = 0.002921), allowing use of Spearman’s correlation test.

**Fig. 3.**
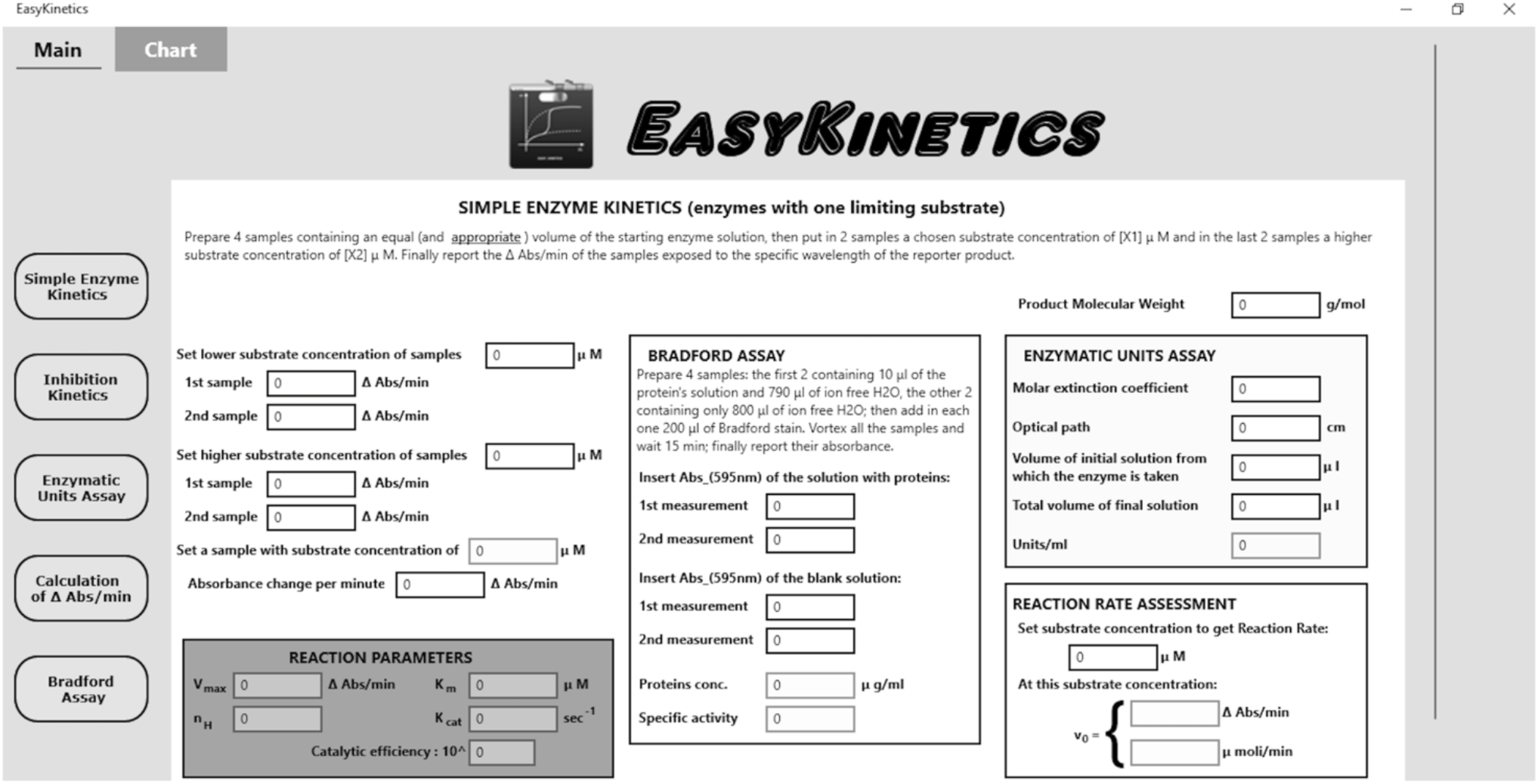
“Simple Enzyme Kinetics” environment: the input fields are characterized by a white background while the output fields by a colored one. On the left are listed the 5 software’s environments while the top’s buttons switch to and from the charts created using the parameters evaluated. Below the title of each environment there is a brief guideline to follow to assure the success of the experiment.

### Kinetic parameters generated with Easy Kinetics shows high accuracy and concordance with those ones generated by GraphPad Prism 8

In order to show Easy Kinetics accuracy in the evaluation of the main kinetic parameter, several time series absorbances were acquired for the enzymes aldehyde dehydrogenase and xanthine oxidase at different concentrations of several limiting substrates for the reaction they catalyzed (**Fig.2A**). Then for each enzyme and each substrate the main kinetic parameters *nH*, K_m_ and V_max_ were generated using Easy Kinetics “Simple Enzyme Kinetics” environment and every two points non repeated permutation of the raw data. In this way it was possible to evaluate the precision of the previous parameters generated by the software as (1-C.V.)*100 for K_m_ and *nH*, showing a normal distribution of values generated for all the tested substrates, and as (1-Q.C.V.)*100 for V_max_, showing a non normal distribution of the values generated for 3 of the tested substrates (**Fig.2E**). In summary the parameters generated show very good precision values for V_max_ (∼90%) and *nH* (100%) in all the reaction models, while a little less for K_m_ (∼78%). Afterward the means values of K_m_ and medians values of V_max_ generated by Easy Kinetics for all the tested substrates were correlated with those ones generated by GraphPad Prism 8© [1] using the same input raw data (**Fig.2F-G**). The correlation analysis shows a significant Pearson’s linear correlation coefficient of 0.99 for V_max_ (t = 10.053, df = 2, p-value = 0.00975), with normally distributed values in both Easy Kinetics (W = 0.87392, p-value = 0.3133) and GraphPad Prism 8© (W = 0.88607, p-value = 0.3652) evaluations, and a significant Spearman correlation coefficient of 1 for K_m_ (S = 0, p-value = 0.08333), with non normally distributed values in both Easy Kinetics (W = 0.67175, p-value = 0.005173) and GraphPad Prism 8© (W = 0.65386, p-value = 0.002921) evaluations.

Surprisingly *nH* values generated for all the substrates tested by Easy Kinetics show a standard deviation δ = 0 around a mean of 1 don’t allowing any correlation test with the corresponding *nH* values generated by GraphPad Prism 8 [1]. Thus for every substrate tested it was evaluated the ratios between GraphPad 8 and Easy Kinetics generated *nH* (0.846 for Glyceraldehyde, 1.249 for Hypoxanthine, 1.024 for NAD+ and 1.045 for Xanthine), showing a mean value of 1.041 with a standard error of se = 0.083 of the ratios normal distribution (W = 0.96402, p-value = 0.8042).

## Discussion

The development of Easy Kinetics was driven by the need to have a publicly available graphical interface software, completely dedicated to perform basic enzyme’s kinetic analyses as an alternative to available commercial software. Testing the kinetics of two enzymes for several limiting substrates has shown that the evaluation of the basic kinetic parameters V_max_, K_m_ and *nH*, from which the other parameters could be generated, gives significantly correlated values, or as regards to *nH* almost the same value, both using several two points non repeated permutations in Easy Kinetics as well as the regression based model in GraphPad Prism 8 [1]. In addition it was shown that every permutation for the same substrate gives in output a parameter values with a precision of ∼90% for V_max_, 100% for *nH* and ∼ 78% for K_m_ using Easy Kinetics environment. These results suggest Easy Kinetics could be used as a valid alternative of the most used commercial software for enzyme’s kinetic analyses GraphPad prism 8 [1], with the advantages it could go deep in the kinetic analysis evaluating other important parameters like: K_cat_, catalytic efficiency or the specific activity of the enzyme, and it’s freely available on public repositories. In addition Easy Kinetics original algorithm for the evaluation of K_m_ and V_max_ tries to be an interesting alternative of usually used regression models, known to have several limitations [12].

## Conclusion

Here it was presented a novel intuitive freely available software, completely dedicated to perform enzymatic kinetic characterizations and based on an original algorithm alternative to the common regression models used by commercial software to evaluate the main kinetic parameters of an enzyme. Requiring less input information than other commercial software like GraphPad Prism 8 [1], Easy Kinetics gives the chance to save time and money during enzyme’s characterization experiments, in addition it allows researchers to go deep in this characterization evaluating several important parameters extremely useful to biochemists. However there are still computational improvements and graphical features achievable that are under development and will be reached in next software releases.

## Acknowledgments

The author declares no conflicts of interests. No funding from any public or private organizations has been used to perform this research. Enzyme’s Kinetics raw data were measured in independent tests inside the Biochemical Department of the University of Pisa. Both the source code and the compiled software are available freely for any user on GitHub. All measures, manipulations and exclusions of the data were reported. Sample size was determined before any data analysis.

## Supplementary tables/figures

### Kinetics equations used from literature

Following it will be reported the main equations used by Easy Kinetics during the generation of several kinetic parameters once evaluated V_max_ and K_m_ values [7-10]:

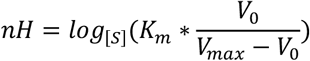

Equation used for the evaluation of the Hill coefficient

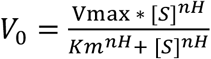

Equation used for the generation of the kinetic graph

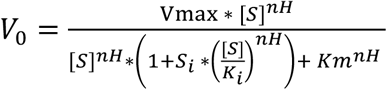

Equation used for the evaluation of the V_0_ at a set chosen substrate

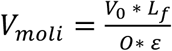

Equation used to switch the previously evaluated V_0_, expressed in ΔAbs/min, into a new V_0_value expressed in μmoli of reporter product generated per minute

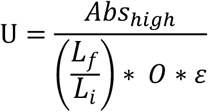

Equation used for the evaluation of the enzymatic units in the sample

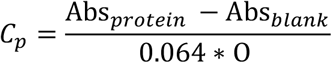

Equation used for the evaluation of the protein concentration during the Bradford assay

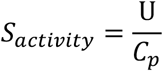

Equation used for the evaluation of the enzyme’s specific activity

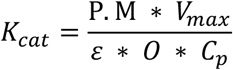

Equation used for the evaluation of the enzyme’s K_cat_

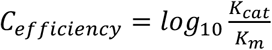

Equation used for the evaluation of the enzyme’s catalytic efficiency

where [S] represents the substrate’s concentration; S_i_ can be 1, if substrate’s inhibition is present or 0, if substrate’s inhibition is absent; K_i_ represents the inhibition’s constant evaluated at a very high substrate’s concentration as:

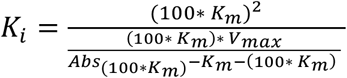

when substrate inhibition is present

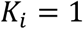

when substrate inhibition is absent

*L*_*f*_ represents the final volume of the sample; *L*_*i*_ represents the starting volume of the sample; ε represents the extinction molar coefficient of the product; O represents the optical path of the spectrophotometer; Abs_high_ represents the absorbance measured at a very high substrate’s concentration; Abs_protein_ represents the absorbance of the protein’s solution; Abs_blank_represents the absorbance measured for the previous solution without proteins inside; P.M. represents the molecular weight of the reporter product.

### Enzyme’s ΔAbs/min raw data for several concentrations of tested limiting substrates

**Tab. 1.** Experimentally measured ΔAbs/min values for several substrate’s concentrations in the enzyme’s catalyzed reactions tested.

| Xanthine oxidase |  |  |  | Aldehyde dehydrogenase |  |  |  |
| --- | --- | --- | --- | --- | --- | --- | --- |
| Hypoxanthine |  | Xanthine |  | Glyceraldehyde |  | NAD+ |  |
| S [uM] | $\Delta Abs/min$ | S [uM] | $\Delta Abs/min$ | S [uM] | $\Delta Abs/min$ | S [uM] | $\Delta Abs/min$ |
| 7,5 | 0,0209 | 5 | 0,0298 | 100 | 0,006 | 10 | 0,008 |
| 10 | 0,0232 | 10 | 0,0412 | 150 | 0,008 | 15 | 0,009 |
| 15 | 0,0300 | 16 | 0,0474 | 200 | 0,009 | 20 | 0,012 |
| 20 | 0,0340 | 20 | 0,0512 | 400 | 0,0125 | 30 | 0,0145 |
| 25 | 0,0366 | 25 | 0,0526 | 600 | 0,0155 | 40 | 0,0165 |
| 30 | 0,0379 | 33 | 0,0529 | 800 | 0,0170 | 60 | 0,0180 |
| 35 | 0,0389 | 40 | 0,0575 | 1000 | 0,0175 | 80 | 0,0195 |
|  |  |  |  | 3000 | 0,0210 | 100 | 0,0205 |
|  |  |  |  | 5000 | 0,0245 | 200 | 0,0230 |

